# Sequential dynamics of Stearoyl-CoA Desaturase-1(SCD1) /ligand binding and unbinding mechanism: A computational study

**DOI:** 10.1101/2020.11.09.373951

**Authors:** Anna B. Petroff, Rebecca L. Weir, Charles R. Yates, Joseph D. Ng, Jerome Baudry

## Abstract

Stearoyl-CoA desaturase-1 (SCD1 or delta-9 desaturase, D9D) is a key metabolic protein that modulates cellular inflammation and stress, but overactivity of SCD1 is associated with diseases including cancer and metabolic syndrome. This transmembrane endoplasmic reticulum protein converts saturated fatty acids into monounsaturated fatty acids, primarily stearoyl-CoA into oleoyl-CoA, which are critical products for energy metabolism and membrane composition. The present computational molecular dynamics study characterizes the molecular dynamics of SCD1 with substrate, product, and as apoprotein. The modeling of SCD1:fatty acid interactions suggests that 1) SCD1:CoA moiety interactions open the substrate binding tunnel, 2) SCD1 stabilizes a substrate conformation favorable for desaturation, and 3) SCD1:product interactions result in an opening of the tunnel, possibly allowing product exit into the surrounding membrane. Together, these results describe a highly dynamic series of SCD1 conformations resulting from the enzyme:cofactor:substrate interplay that inform drug-discovery efforts.

## Introduction

Stearoyl-CoA desaturase-1 (SCD1), an endoplasmic reticulum membrane enzyme, is a central regulator of energy metabolism[1]. SCD1 desaturates stearoyl-CoA and palmitoyl-CoA, into monounsaturated fatty acids (MUFA) oleoyl-CoA and palmitoleoyl-CoA through the insertion of a double bond in the Δ-9 position of the substrate[2], as indicated by SCD1’s alternate name, delta-9 desaturase (D9D). This oxidative reaction requires electron transport cytochrome b_5_ and molecular oxygen[2] (Fig 1, A - C). Well-controlled activity of SCD1 is critically important to health as the products are used in the formation of phospholipids, triglycerides, and cholesteryl esters, and hence contribute to membrane fluidity, adiposity, and signal transduction[3]. SCD1 is ubiquitously expressed, particularly in lipogenic tissues, and its concentration changes in response to hormonal and dietary effectors[4].

**Fig 1.**
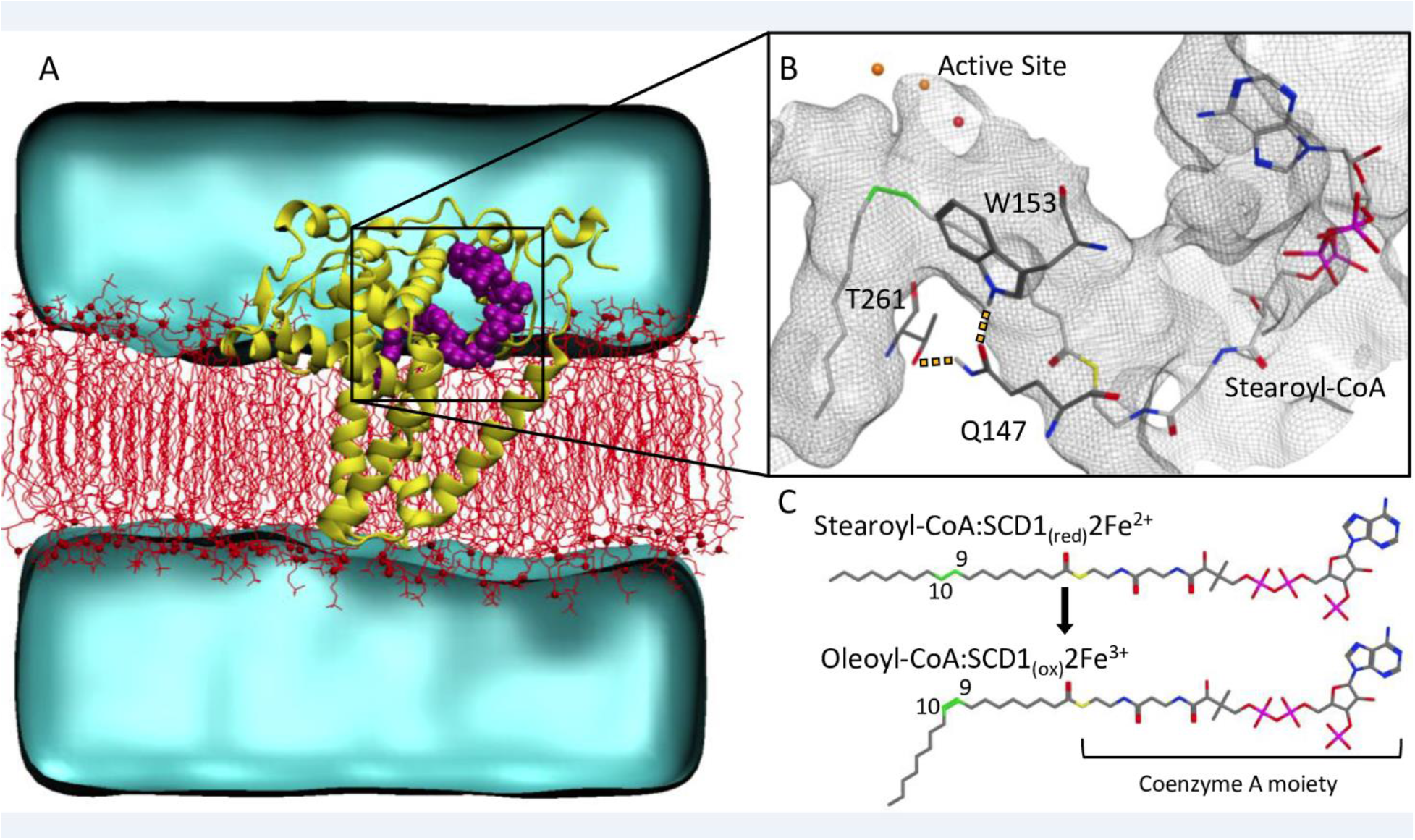
Stearoyl-CoA Desaturase 1 Δ-9 Desaturation Reaction. (A) SCD1 desaturase (yellow) in POPC membrane (lipids as red lines, phosphate groups as red spheres); with stearoyl-CoA substrate (purple); water environment (cyan). (B) SCD1 hydrophobic tunnel (gray mesh) with stearoyl-CoA (colored by atom type). Substrate 9^th^ and 10^th^ carbons (green) positioned proximal to the active site’s diiron center (orange) and oxygen atom of crystallographically-determined water (red). Hydrogen bonding between residues in yellow dashes. (C) The SCD1 desaturation reaction, resulting in a double bond at the Δ-9 position, converting stearoyl-CoA (substrate) to oleoyl-CoA (product).

While SCD1 activity is critical to health, particularly because it is important to the modulation of cellular inflammation and stress[5], SCD1 activity can be associated with disease under certain conditions. Under high-fat conditions, SCD1 deficiency protects mice from obesity and insulin resistance[6] and hepatic steatosis[7]. Similarly, high SCD1 activity predicted the development of metabolic syndrome in men[8]. SCD1 overexpression is implicated in cancer[9,10] and it has been described as an oncogene[11]. As such, SCD1 is a potentially important pharmaceutical target: inhibitors of SCD1 have been developed as potential treatments for diabetes[12] and cancer[13,14]. Because SCD1 activity has nutritional effectors, nutraceuticals have been investigated as mitigators of SCD1 overactivity. For example, Manni *et al*.[15] found that treatment with the drug Lovaza®, made of ethyl esters of the omega-3 fatty acids DHA and EPA, resulted in lower levels of SCD1 products in obese postmenopausal women at high risk for breast cancer. In addition, results from a mouse model of alcoholic liver disease indicate that the naturally occurring SCD1 inhibitor *Lycium barbarum* polysaccharide, has liver-protective effects[16]. Nonetheless, the development of pharmaceutical or dietary modulators of SCD1 has been challenging because of variance in its expression by organ type[17].

The recent determination of human[18] and murine[19] SCD1 crystal structures provides fundamental insights into the mechanism of action of the enzyme and of its specificity. These findings open the door to a rational, structure-based modulation of its activity. Both human and murine SCD1 structures exhibit a bent, or kinked, binding site for the saturated-CoA substrate. In their murine model, Bai *et al*.[19] attribute the kink to the hydrogen-bond interactions between three residues that underlie the binding tunnel. The homologous human protein structure[18] reveals the same interaction pattern between these conserved residues: Gln147 interacts with both Trp153 and Thr261 (Fig 1, B). Bai *et al*.[19] suggest that this binding site geometry drives the conformation of the saturated substrate acyl chain such that the regioselective desaturation can occur at C-9, as was first hypothesized in 1969[20]. Inhibition and analog studies[2] indicate that the carbons C-9 and C-10 of the substrate’s lipid tail adopt a gauche conformation in order for the rate-limiting hydrogen abstraction that results in the *cis* double bond to occur. However, the crystal structure of human SCD1 exhibits a non-gauche conformation with a −111.1° C8-C9-C10-C11 dihedral angle[18]. The human and mouse SCD1 crystal structures also exhibit several non-bonded contacts between the CoA moiety of the substrate and the protein’s cytoplasmic domains, suggesting a possible active effect of CoA in the positioning of substrates and consistent with assay results indicating that the CoA moiety is required for binding[2].

SCD1’s human disease relevance, including cancer and metabolic syndrome, makes it is important candidate disease target. However, although there are numerous studies concerning *scd1* expression, genetic variation, and transcriptional regulation[3–5,7,21–23], the only known mutagenesis studies on the SCD1 protein concern either the residues that contribute to catalysis[24] or the residues that may be mutated to prevent the quick degradation of this short-lived membrane protein[25]. Mutagenesis results from other desaturases, such as those that desaturate a different carbon-carbon bond (e.g., delta-6 rather than delta-9), interact with a different headgroup (e.g., Acyl Carrier Protein rather than CoA), or occupy a different area of the cell (e.g., solvated vs. membrane), do not provide results that can be extrapolated to inform the workings of SCD1. For instance, rat delta-6 desaturase[26] is also a membrane protein that desaturates lipids with CoA headgroups. While mutagenesis studies indicate that the binding tunnel has residues that contribute to substrate selection[26], the rat D6D-human D9D identity is only 21%. Furthermore, it is not known if rat D6D has a similar structure to human D9D despite this difference in identity. Vanhercke et al.[27], who published mutagenesis results on a bifunctional Δ-12/Δ-9 membrane-bound desaturase found in crickets, state that due to lack of structural data on membrane-proteins “our understanding of the structure-function relationship of membrane-bound desaturases remains limited and scattered at best” (pg.12860). Similar efforts to characterize catalysis on fatty acids face the same problem: even when site-directed mutagenesis studies exist, without structural data, interpretation is limited[28]. The recent structures of SCD1[18,19] open the door to a much-needed functional characterization of the ligand:protein interactions beyond the catalytic mechanism.

A better understanding of the substrate-protein interaction in the binding site dynamics of SCD1 and its substrates is needed. Hence, these results help to guide and prioritize future mutagenesis wet-lab experiments. The recently determined crystal structures identified three residues thought to confer a kinked binding tunnel that facilitates substrate position for desaturation. However, it is unknown to what extent the binding tunnel changes shape across the protein turnover cycle (from apoprotein to protein-substrate to protein-product, beginning again as apoprotein). In this study, we characterize SCD1’s conformational states by *in silico* dynamic modeling to identify how the distances between the three residues of interest correspond to these different states of the protein turnover cycle and to inform how the CoA moiety contributes to these differences.

## Materials and methods

### Model construction and simulations

All models (described in Table 1) are based on PDB entry 4ZYO, the human SCD1 structure determined by Wang *et al*.[18]. In building the transmembrane SCD1 model, enzyme and substrate atoms were included in the model. The co-crystalized dodecyl-beta-D-maltoside was deleted and the zinc ions in the crystal structures were replaced by the naturally occurring iron ions at the same locations. The force field parameters for stearoyl-CoA were generated with CHARMM General Force Field (CGenFF)[29] and using the CHARMM36[30] force field for the protein, lipids, water, and ions parts of the model. Membrane generation, placement, and preparatory steps were performed using the QwikMD[31] and NAMD2[32] programs. From an initial orientation obtained using the OPM server[33], the protein was placed into a model membrane made of 1-palmitoyl-2-oleoyl-*sn*-glycero-3-phosphocholine (POPC), the most common ER membrane lipid species[34]. The protein and membrane were solvated with TIP3P water models and with salt concentration of 0.15mol/L of NaCl in a periodic box of 100 Å x 100Å x 100 Å, representing a total of 95,297 atoms.

**Table 1.**
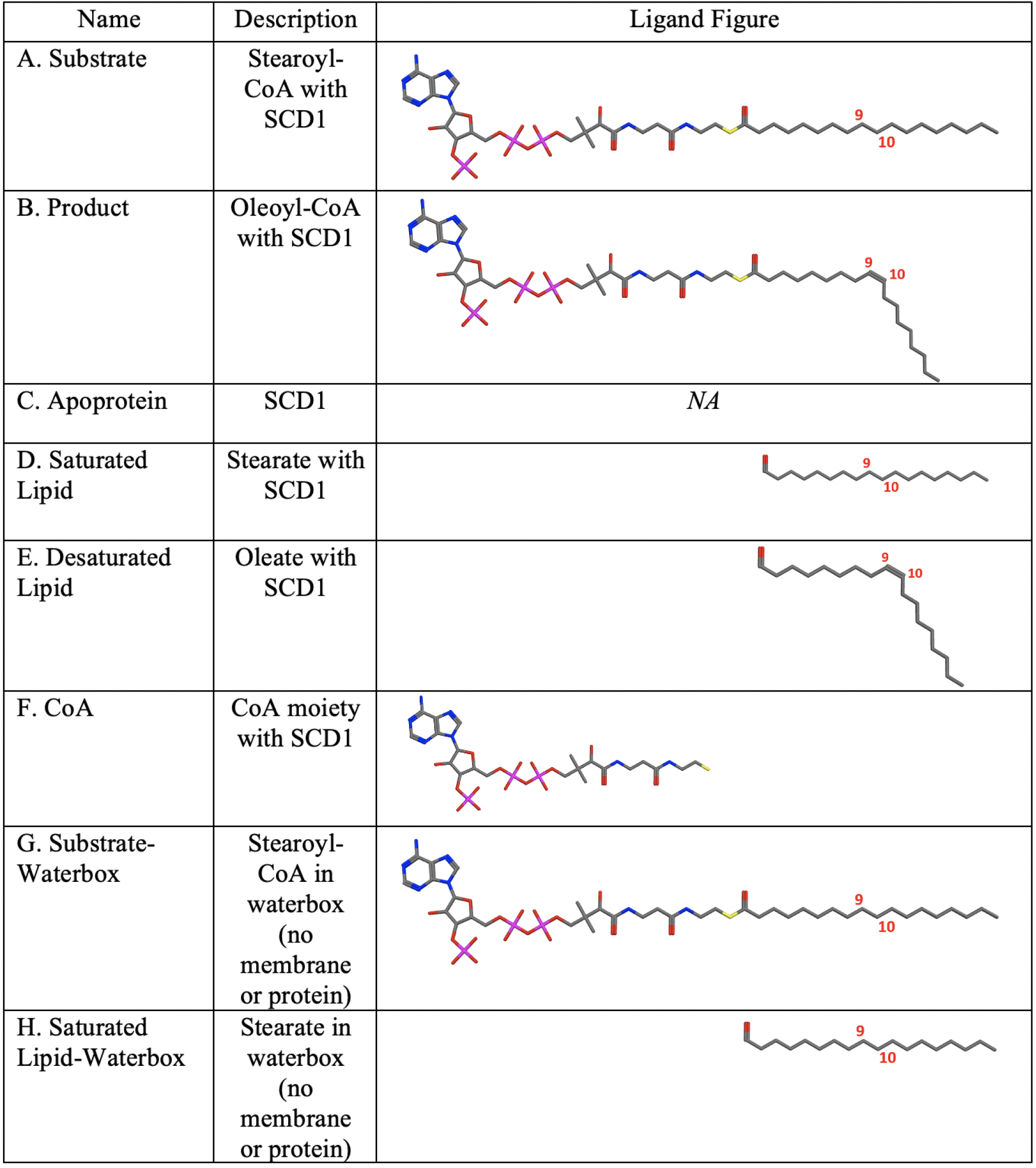
Model names and descriptions.

| Name | Description | Ligand Figure |
| --- | --- | --- |
| A. Substrate | Stearoyl-CoA with SCD1 |  |
| B. Product | Oleoyl-CoA with SCD1 |  |
| C. Apoprotein | SCD1 | *NA* |
| D. Saturated Lipid | Stearate with SCD1 |  |
| E. Desaturated Lipid | Oleate with SCD1 |  |
| F. CoA | CoA moiety with SCD1 |  |
| G. Substrate-Waterbox | Stearoyl-CoA in waterbox (no membrane or protein) |  |
| H. Saturated Lipid-Waterbox | Stearate in waterbox (no membrane or protein) |  |

The membrane-protein model was submitted to the following procedure, all using explicit hydration, with a 12Å non-bonded interactions cutoff, and using Particle Mesh Ewald. First, the heavy atoms of the protein backbone, the active site residues (HIS120, HIS125, HIS127, HIS160, HIS161, ASN265, HIS269, HIS298, HIS301, HIS302), and the stearoyl-CoA ligand were restrained using a harmonic restraint of 2 kcal/mol Å^2^ about their crystallographic positions. Next, the membrane lipids were annealed to the protein (i.e., allowed to surround the protein as they would in a biological membrane) over the course of 30 ns with the same active site restraints using the QwikMD Toolkit in VMD. The model system was then equilibrated for 1 ns with the same restraints on the atomic positions. Finally, an additional 1-ns molecular dynamics (MD) simulation equilibration was run with the stearoyl-CoA acyl tail carbons restrained and the CoA unrestrained. The resultant model system contained no water molecules inside the membrane. The equilibrated model is shown in Fig 1-A. As shown, the protein structure was oriented such that the four transmembrane helices extend across the membrane, with amphipathic helices at the membrane-solvent interface and protein cap with the active site extending into the cytosol (Fig 1-A). This position is consistent with that described in Wang *et al*.[18]. The average membrane thickness around the protein during the MD simulation was 32.1 +/− 0.2 Å.

Using this equilibrated model, a series of different models were built, with each differing from each other by the type of ligand present. These modified ligands, shown in Table 1, were constructed using the program MOE version 2019 (Molecular Operating Environment; Chemical Computing Group, Montreal, Canada), which generates Amber 99 force field parameters for each ligand. For each variant, the solvent was first deleted from the membrane-protein system. The desired ligand was included in the model, and the system was re-solvated using the MOE solvate facility 0.15 mol/L Na^+^Cl^−^ counterions. The distances between the iron ions and the coordinating histidine or water residues were restrained around their crystallographic positions using harmonic potentials of 40 kcal/mol Å^2^. Each model was included in a 100 Å x 100Å x 100 Å periodic box, which totaled between 99,577 and 100,161 atoms, depending on the model. The models are: 1) Model “Substrate”: stearoyl-CoA, i.e., with a lipid fatty acid chain with a C9-C10 single bond, (Fig 1-A) in SCD1, as shown in Table 1-A. 2) Model “Product”: oleoyl-CoA, i.e., with a lipid fatty acid chain with a C9-C10 double bond, in SCD1, as shown in Table 1-B. 3) Model “Apoprotein”: SCD1 without substrate, as in in Table 1-C. 4) Model “Saturated Lipid”: Stearate (i.e., fatty acid as in Table 1-A without the CoA moiety) with SCD1, as shown in Table 1-D. 5) “Desaturated Lipid”: oleate (i.e., fatty acid as in Table 1-B without the CoA moiety) in SCD1, as shown in Table 1-E; 6) “CoA”: CoA moiety, without the fatty acid, in SCD1, as shown in Table 1-F. In order to account for the protein tunnel’s contribution to ligand shape, two additional models were built containing only ligands in a waterbox (i.e., containing no protein and no membrane): 7) “Substrate-waterbox”: stearoyl-CoA (i.e., as in Table 1-A) in a periodic water box 40 Å x 40Å x 40 Å, as shown in Table 1-G. 8) “Saturated Lipid-waterbox,”: stearate (i.e., as in Table 1-D) in a periodic water box 40 Å x 40Å x 40 Å, as shown in Table 1-H.

Each model was energy-minimized to an energy gradient of less than 10^−4^ kcal/mol/Å^2^ followed by a 1-ns MD equilibration simulation. Subsequently, a 100-ns production run was performed for a total combined production simulation time (for all eight models) of 800ns.

### Molecular dynamics analysis

MD trajectory measurements were used to measure the dihedral angle values of C-8, C-9, C-10, and C-11, as well as the distance between the two irons of the active site and the distance from each iron to C-9 and C-10. Both human and murine crystal structures indicated a kinked binding tunnel. Bai *et al*.[19] proposed that the tunnel kink is due to hydrogen bonding between Gln147, Trp153, and Thr261. The present work defined these distances as follows: 1) distance between Gln147 (Oɛ1) and Trp153 (hydrogen of indole nitrogen Hɛ1) and 2) distance between Gln147 (H ɛ22) and Thr261 (Oγ1) as shown in Fig 1-B. In the human crystal structure[18], the 5th carbon of the stearoyl-CoA lipid tail appears at the tunnel entrance. The overall structure was solved at 3.25 Å resolution. Several residues close to the residues of interest listed here exhibited relatively high B factors: Trp153 (atoms C, CD2, and CE3, with B factors of 101.69, 101.14, and 100.12, respectively), and Thr143 (atom OG1 with B factor of 106.69). Volume measurements were centered on this 5th carbon and extend at a 6Å radius, resulting in a sphere that includes the three residues of interest. Volume measurements were obtained using the program POVME3[35] from the 100-ns MD trajectories.

## Results

### Structure and dynamics of Substrate, Product, and Apoprotein models

Comparison of the Substrate, Product, and Apoprotein models (Table 1-A, 1-B, and 1-C, respectively), indicates that each of these models has a characteristic mean distance between the Gln147, Trp153, and Thr261 residues (SI Table 1) suggested to be responsible for the kink conformation of the binding site[19]. SI Figs 1 and 2 show the fluctuation of these distances across the 100-ns trajectory for each model. When these distances are grouped into 10-ns bins (i.e. first bin includes the first 10ns; SI Fig 3) the Substrate model (Table 1-A) distances remain consistent across the 100-ns trajectory (SI Figs 3A and 3B), whereas the Product model distances within the first 10-ns bins differ from those of the remaining 90-ns (SI Figs 3C and 3D). This difference in the Product model (Table 1-B) is arguably artifactual, as the crystal structure of D9D used as the basis for the Product model (Table 1- B) was co-crystalized with the substrate rather than product ligand. In order to account for the effect of this difference, the first 10-ns of the Substrate and Product model (Table 1-A, B, respectively) trajectories was omitted from the following analyses.

### Substrate tunnel kink and protein’s hydrogen bond network

For each model, every observed paired distance between Gln147-Trp153 and Gln147-Thr261 was grouped into bins of tenths of angstroms, (distribution by population for all models in SI Fig 4). The free energy difference between these conformations, ΔG, for the Substrate, Product, and Apoprotein models is shown in Fig 2-A, B, C, respectively (Table 1-A, B, respectively). ΔG was calculated as -kT ln(ρi/ρmax), with ρ_i_ the population of bin i, and ρ_max_ the population of the most populated bin. Bins without observations were given an arbitrary ΔG value of 8 kT, i.e., higher that corresponding to the less populated bin.

**Fig 2.**
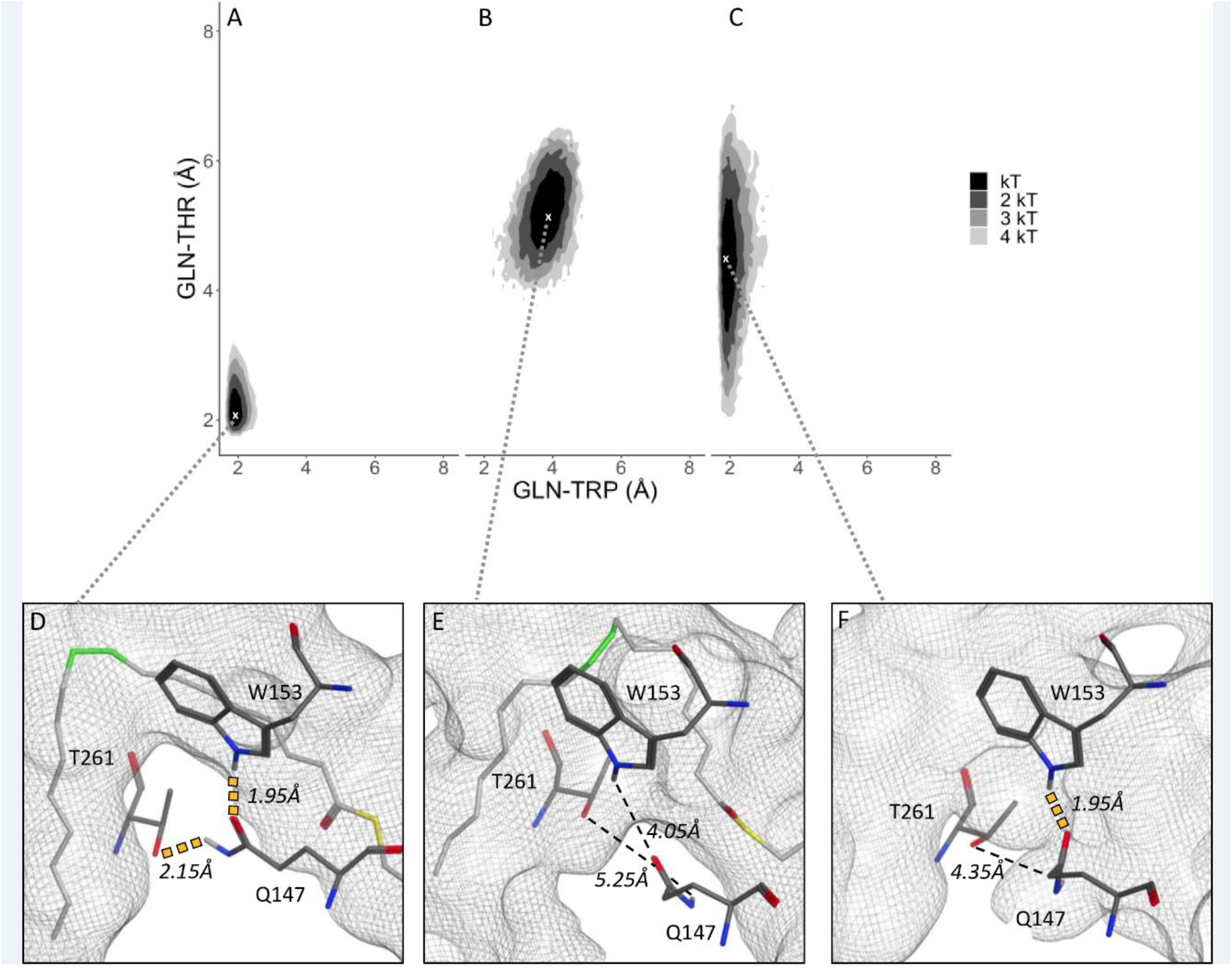
ΔG distribution of the paired Gln-Trp and Gln-Thr distances. (A) Substrate; (B) Product; (C) Apoprotein models. Minima indicated with white “x”. D, E, F: Binding tunnel (gray mesh) for each corresponding ΔG minima of A, B, and C, with distances between Gln-Trp and Gln-Thr within hydrogen bonding range (yellow dotted line) and outside of hydrogen bonding range (black dashed line).

For the Substrate model (Table 1-A), the ΔG minimum was at Gln147-Trp153 = 1.95Å and Gln147-Thr261 = 2.15Å and included 4.7% of total observations, whereas the Product model (Table 1-B) had a ΔG minimum of Gln147-Trp153 = 4.05Å and Gln147-Thr261 = 5.25Å and included 1.0% of total observations. While the Substrate and Product distributions overlapped, there were no Substrate observations at the Product distribution minimum or vice versa. The Apoprotein model (Table 1-C) displayed a ΔG minimum at Gln147-Trp153 = 1.95Å, whereas for Gln147-Thr261 the ΔG minimum extended across two bins with same population. This finding indicates that the most observations extended not within one tenth of an angstrom, but within two, varying from 4.3Å to 4.5Å and including a total of 2.3% of observations. Table 2 presents the percent of observations at each group in order of increasing ΔG kT levels.

**Table 2.** Percentage of paired Gln147-Thr261 and Gln147-Trp153 distances for each model grouped by kT.

|  | Substrate | Product | Apoprotein | Sat.Lipid | Desat.Lipid | CoA |
| --- | --- | --- | --- | --- | --- | --- |
| Minima | 4.7 | 1.0 | 2.3 | 2.7 | 1.1 | 0.6 |
| 1 kT: <0.6 | 54.9 | 57.2 | 50.5 | 56.1 | 18.4 | 46.6 |
| 2 kT: 0.6-1.19 | 23.5 | 25.4 | 25.7 | 24.0 | 21.2 | 34.2 |
| 3 kT: 1.2-1.79 | 9.5 | 9.2 | 12.2 | 7.8 | 29.9 | 10.9 |
| 4 kT: 1.8-.2.39 | 4.5 | 3.8 | 5.8 | 4.3 | 16.4 | 6.0 |
| 5 kT: 2.4-2.99 | 1.8 | 2.0 | 2.0 | 2.5 | 8.8 | 1.8 |
| 6 kT: 3.0-3.59 | 0.6 | 1.3 | 1.4 | 1.5 | 4.2 | 0.0 |
| 7 kT: 3.6-4.19 | 0.4 | 0.0 | 0.0 | 1.0 | 0.0 | 0.0 |

The geometry of the binding sites corresponding to the minimum free energies of the three models are shown on Fig 2- D, E, & F. In the Substrate model (Table 1-A), the gray mesh surface of the tunnel is in the crystallographic kinked geometry (Fig 2-D), whereas there is no such kink in the Product model (Table 1-B), as shown by the comparatively more spacious gray mesh tunnel (Fig 2-E). Taken together, the three residues are closer together in the Substrate model (Table 1-A) and are observed within a kinked tunnel, whereas the three residues are farther apart in the Product model (Table 1-B), which is not kinked. The distribution of Gln147-Thr261 and Gln147-Trp153 distances in the Apoprotein model (Table 1-C) exhibits distributions that are in between those of the Substrate and Product. The trajectory of these distances, shown in SI Figs 1 and 2, respectively, indicates that the Gln147-Trp153 distance remained more consistently close compared to the Gln147-Thr261 distance, which fluctuated. Although these residues are distinct in the Substrate vs. Product conformations, the distances between the two active site irons and the C-9 and C-10 carbons is similar in the two models. The respective distance for the Substrate and Product models, in comparison, are as follows: Iron 1 to C-9, 7.5 Å and 7.2 Å, respectively, Iron 1 to C-10, 7.1 Å and 6.9 Å, respectively, Iron 2 to C-9, 6.8 Å and 7.1 Å, respectively, and Iron 2 to C-10, 6.9 Å and 6.8 Å, respectively. The distance between the two active site irons was also similar in the Substrate and Product models, 7.4 Å and 8.7 Å, respectively. Findings from the present work suggest a highly dynamic role of the three residues of interest, specifically one that varies in relation to the absence of a ligand, the presence of saturated substrate, or the presence of a desaturated product.

### Substrate tunnel kink and C9-C10 dihedral

The crystallographically-observed kink of the Substrate model (Table 1-A) contributes to a favorable positioning of the substrate in the tunnel. SCD1 desaturates at C9-C10 of the lipid tail of the stearoyl-CoA substrate. A negative gauche dihedral about C9-C10, rather than a transconformation, was proposed as the favored conformation for the desaturation reaction[2]. Fig 3-A shows ρ(χ), the distribution of the χ dihedral angle of the stearoyl-CoA formed by the 8^th^,9^th^,10^th^ and 11th carbon atoms in the “Substrate” model (Table 1-A), “Substrate-Waterbox” model (Table 1-G), and “Saturated Lipid-Waterbox model” (Table 1-H). The free energy corresponding to rotations around this dihedral, calculated as ϕ = −0.6 ln (ρ(χ)) (in kcal/mol), is represented by the overlaid gray line. The corresponding value of this dihedral angle in the crystal structure, −111.1°, is indicated with an asterisk in the bottom plot. The dihedral distribution observed in the “Substrate” model, i.e., with the saturated C9-C10 bond in the protein, shows a nearly-equal population of trans and of negative gauche dihedral conformations. The Substrate-waterbox distribution indicates a preferred trans C9-C10 dihedral in solution (around 180°) for the substrate, with a negative gauche C9-C10 dihedral (−60°) which is about 7.5 times less populated than the trans conformation, and with a free energy barrier separating these two conformations of about 4 kcal/mol (Fig 3). Comparison of these models suggests that the protein environment of the substrate catalyzes the population of an actionable conformation for the C9-C10 bond in the substrate from mostly trans (in solution) to equi-probable cis and trans in the protein In order to investigate the role of the CoA moiety in attaining a gauche C9-C10 dihedral, these findings were compared to those of the Saturated Lipid Model (Table 1-D), which showed a strongly favored negative gauche conformation (around −60°), which is in this case 6.6 times more populated than the trans conformation. This finding suggests that the CoA headgroup hinders the formation of conformations compatible with desaturation, and that, on the other hand, the interaction between the lipid tail and the protein tunnel promotes conformations compatible with desaturation.

**Fig 3.**
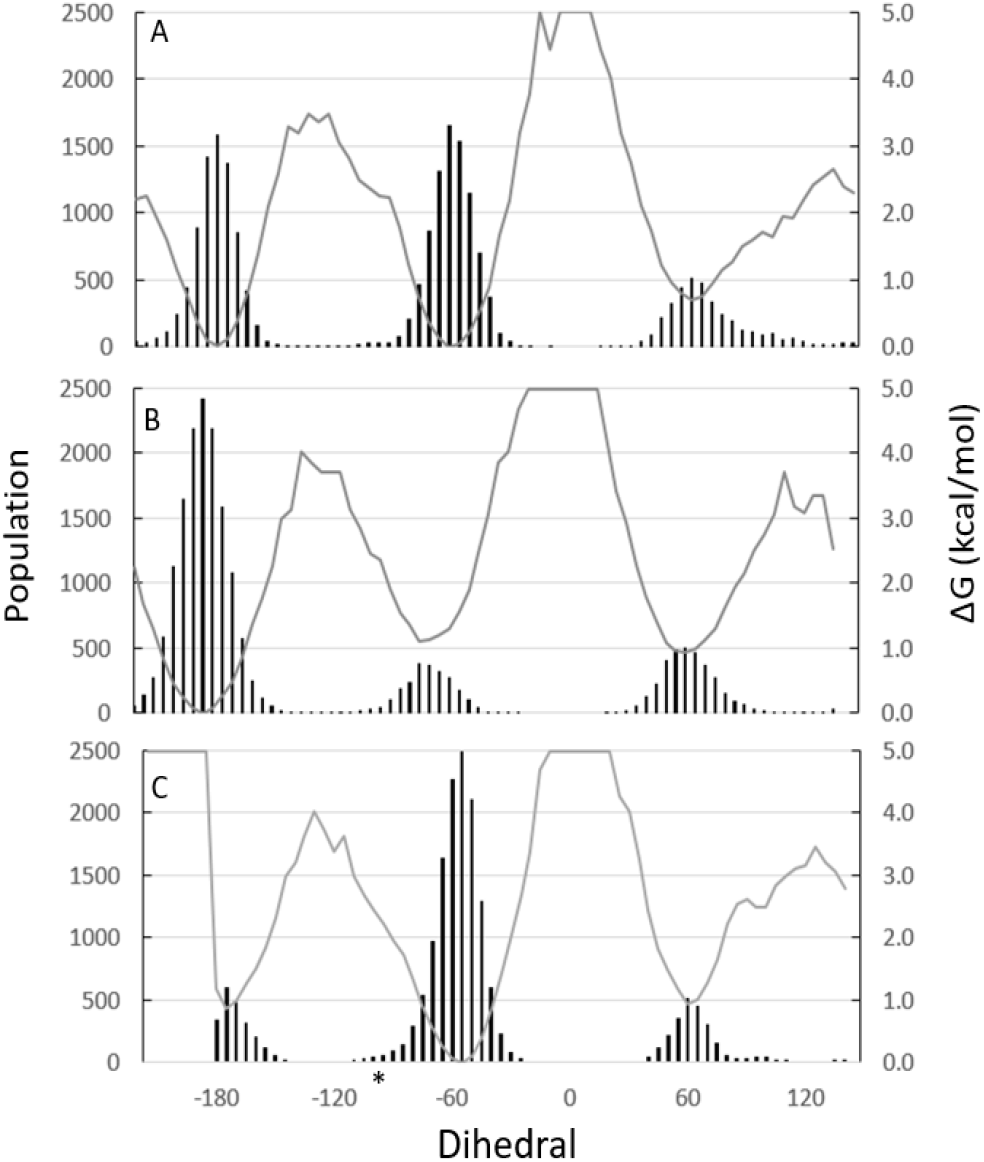
Distribution of ligand C9-C10 dihedral populations. MD populations of dihedral angles (black histogram lines) with corresponding ΔG (gray line). (A) Substrate-Protein; (B) Substrate-waterbox; (C) Saturated Lipid-Protein. The crystallographic dihedral value (−111.1°) is indicated by an asterisk in panel C.

### Role of CoA in the substrate conformation

Comparison of the Apoprotein model (Table 1-C) and the CoA model (Table 1-F) shows interactions between the protein and CoA moiety may contribute to substrate entry into the protein. The three residues of interest, Gln147, Trp153 and Thr261, are located at the tunnel entrance (Fig 4). Comparison of the free energy distributions based on these distances for the Apoprotein and CoA models shows that while the Apoprotein model fluctuated between resembling the Substrate and Product models, the CoA model exhibited a markedly greater distance between these three residues, particularly the distance between Gln147-Thr261. The distance between Gln147-Thr261 is significant for substrate entry because these residues are across from each other underneath the binding tunnel, and as such, separation in this area may contribute to entry of the substrate from the surrounding membrane (Fig 1-A, B). When the CoA moiety interacted with the protein face, as represented by the CoA model (Table 1-F), there was a substantial population of observations with a distance of Gln-Thr 8.25 Å (ΔG less than 0.6 kcal/mol), which is 1.8x greater than that of the Apoprotein model (Table 1-C).

**Fig 4.**
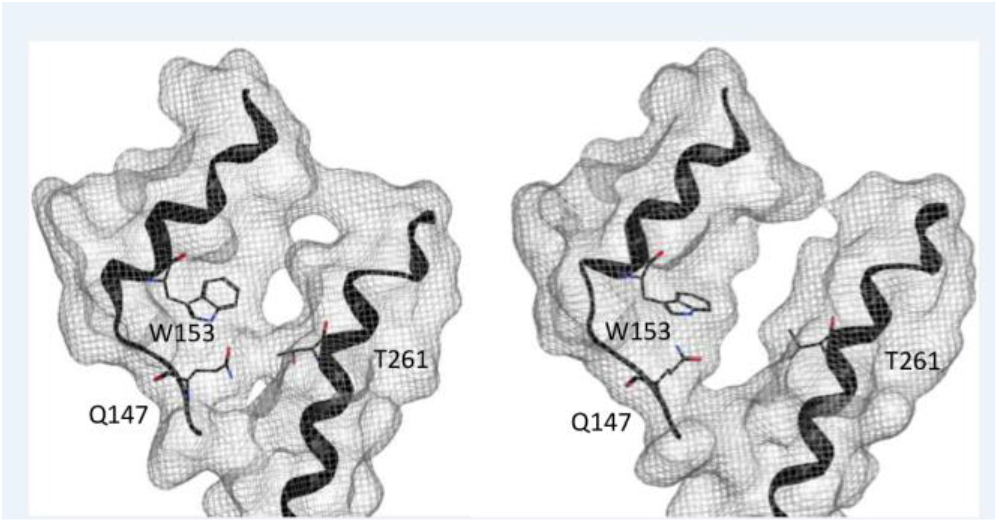
Structure of the enzyme’s tunnel entrance. Apoprotein model (left) and CoA model (right) with tunnel surface in gray mesh.

Gln147 is located on a short loop between Trp153 and Thr261 (SI Fig 5, inset). In the Apoprotein model (Table 1-C), this loop remained in a fairly stable conformation (SI Fig 5) and maintained the Gln147-Trp153 distance while allowing some fluctuation of the Gln147-Thr261 distance. The volume at the protein interface (shown in SI Fig 6) remained relatively stable, fluctuating between 75.2Å3 and 78.4Å3 (SI Fig 6). In the CoA model (Table 1-F), the loop including Gln147 exhibited a conformational change 27.5ns into the MD trajectory (SI Fig 5). Following this change, the distance between Gln147 and Thr261increased (SI Fig 1-f). Consistent with this greater distance, the volume of the tunnel entrance increased (SI Fig 6) from 51.4Å3 to 136.9Å^3^. The tunnel entrance is visibly greater, as shown in Fig 4. Fig 5 presents the tunnel entrance volume, which shows that the Apoprotein model tunnel entrance was partially kinked (banana-shape) and the CoA model (Table 1-F) tunnel was not kinked (open). Taken together, the interaction of the CoA moiety with SCD1 results in a greater distance between Gln147 relative to Thr261, an increase that corresponds to in an increased volume at the tunnel entrance. In this way, the CoA moiety may aid in substrate recognition and entry into the binding tunnel.

**Fig 5.**
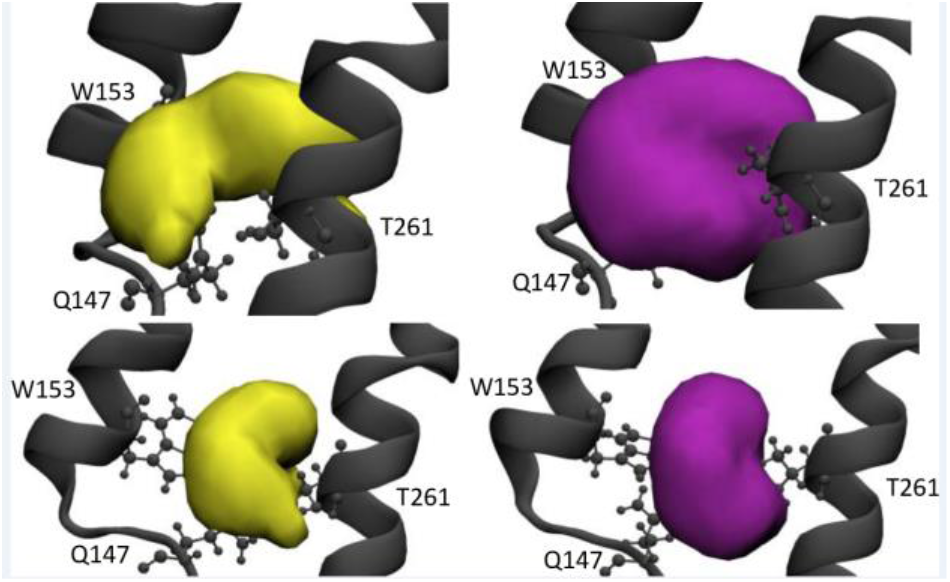
Tunnel entrance volume comparison. Unoccupied tunnel entrance volume in the Apoprotein (gold) and the CoA (purple) models. The bottom panel is rotated 60° to the left respective to the top panel.

### Discrimination between substrate and product

The results indicate that the CoA headgroup contributes to the protein discrimination between substrate and product, as the presence or absence of the CoA moiety revealed differing effects on the paired residues depending on the saturation or desaturation of the remaining acyl tail. In the case of the Saturated Lipid model (Table 1-D), the loss of the CoA moiety resulted in a slight displacement of the ΔG minimum, namely a 0.3Å increase in the Gln147-Trp153 distance compared to the Substrate Model, an increase reflective of a secondary subpopulation of observations with a greater Gln-Trp distance (SI Fig 4-D). However, for the Desaturated Lipid model, loss of the CoA moiety resulted in a novel minimum (SI Fig 4-E). While the Product model minimum was at Gln147-Trp153 4.05Å, Gln147-Thr261 5.25Å, in the Desaturated Lipid model it was at Gln147-Trp153 1.95Å, Gln147-Thr261 2.25Å (SI Fig 4-B, E, respectively). This novel minimum was similar to the Saturated Lipid model (i.e., a 0.1Å increase in the Gln-Thr distance). Taken together, the CoA moiety has a role in Substrate-Product discrimination, as loss of the CoA moiety reduced the distinction between Substrate and Product ΔG distributions (SI Fig 4 A vs D, B vs. E; Table 1-A,B, respectively).

## Discussion

These results provide insight into the mechanism of SCD1’s interactions, from the apoprotein state to the initial contact of the substrate’s CoA moiety to the positioning of the substrate in the tunnel to the discrimination between substrate and product. Scheme 1 illustrates this proposed series of events: 1) The apoprotein exhibits a shallow but open tunnel entrance 2) CoA-moiety interacts with the protein surface, resulting in an increased tunnel entrance volume. 3) After the substrate inserts in the protein, the tunnel is kinked. 4) After the substrate is catalyzed into the product, the tunnel shape is no longer kinked, possibly allowing for egress of the product into the membrane. 5) Product release indicates the presumed transition from Product model to Apoprotein model, which was not simulated in the present work.

**Scheme 1.**
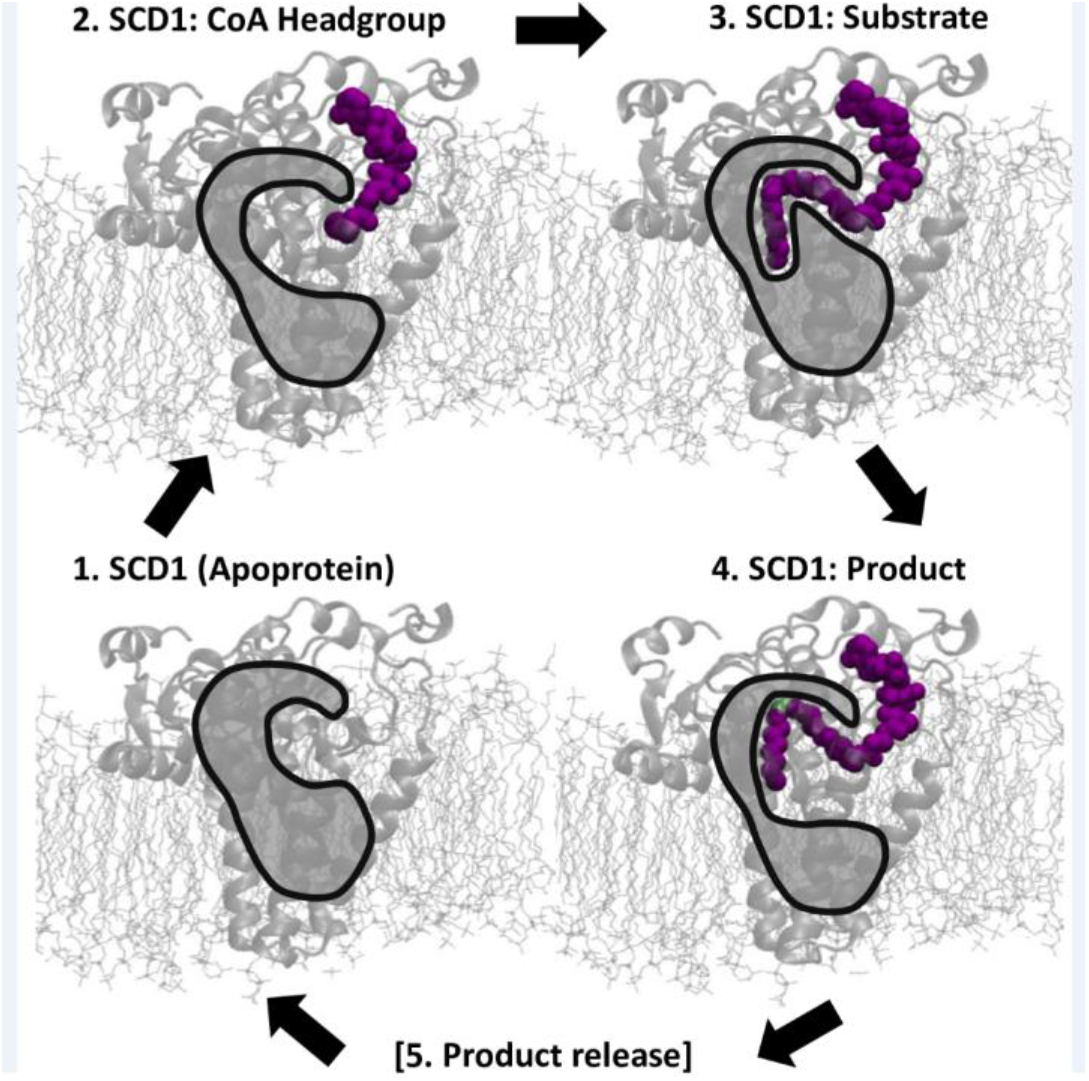
Proposed sequence of binding events. SCD1 in the membrane. The hydrophobic tunnel is represented by an opaque gray shape with black outline with purple ligand, desaturated carbons in green.

### From apoprotein to substrate insertion (Scheme 1, steps 1 and 2)

Based on their crystal structure observations, Wang *et al*.[18] hypothesized that the order of interactions between the stearoyl-CoA substrate and apoprotein begin with the interaction of the CoA moiety and the nine protein face residues, followed by the movement of the fatty acid tail proceeding towards the tunnel entrance, and passing between TM2 and TM4. The present results are largely consistent with this view. The tunnel entrance opening, which is the space between the three residues of interest, is smaller in the Apoprotein model than the CoA model (Table 1-F, 1-C, respectively; Fig 5). As shown in Fig 4, the Apoprotein tunnel entrance is constricted by the three residues, whereas in the CoA model the greater distance between Gln147 and Thr261 allows for a greater entrance opening. Because the increased distance between Gln147 and Thr261 was observed in the CoA model (Table 1-F) but not in the Apoprotein model (Table 1-C), the tunnel-opening action of CoA appears to be due to an allosteric change based on the interaction between these nine residues and the CoA moiety. Results suggest that this interaction of the CoA moiety with the protein (Table 1-F) results in a rearrangement of the loop at the tunnel entrance followed by an 62.5% increase in the tunnel entrance volume (SI Figs 5 and 6, respectively). This finding is consistent with experimental results that indicate that SCD1 acts on stearoyl-CoA but not stearate, which is the saturated lipid tail without the CoA headgroup[2].

Given these findings, the series of actions leading to stearoyl-CoA entry into SCD1 may be as follows (see *Scheme 1, steps 1 and 2*): The residues that the CoA moiety interact with serve as a “doorbell” for the CoA head group of the incoming substrate to press. Following this action, the loop including Gln147 rearranges such that interactions with Thr261 are destabilized, resulting in a greater tunnel entrance volume. While the distance between Gln147-Thr261 grows, the Gln147-Trp153 remains close, possibly guiding the loop movement. These dynamics may facilitate the entrance of the substrate from the membrane into the protein’s hydrophobic tunnel.

Mutagenesis studies of the membrane desaturase rat D6D, which does not presently have a determined structure, indicated eight tunnel residues that contribute to substrate specificity, including a Trp, Gln, and Ser26. As these residues contribute to substrate specificity, they have been proposed to be near the binding tunnel entrance. However, due to the low identity and differing substrates between rat D6D and human D9D, it is not possible to assert whether or not if the rat D6D’s Trp, Gln, and Ser residues are equivalent, structurally and functionally, to the Trp, Gln, and Thr described in the present work on human D9D.

### From inserted substrate to product (Scheme 1, steps 3 and 4)

The Substrate model (Table 1-A) free energy distribution indicates that when the two paired distances, Gln147-Trp153 and Gln147-Thr261, were simultaneously close (1.95Å and 2.15Å, respectively), the dihedral about the stearoyl-CoA C9-C10 bond was actionable (−60° +/− 10°). This finding indicates that for the Substrate model, the lower the free energy of the paired Gln147-Trp153 and Gln147-Thr261 distances, the greater the number of actionable dihedrals: the proportion of actionable dihedrals decreases with the increase from ΔG 1 kT, 2 kT, 3 kT, 4 kT (52.1%, 25.6%, 5.3%, 4.8%, respectively). Taken together, the Gln147-Trp153 and Gln147-Thr261 paired distances are significantly related to the favorable positioning of the substrate. As shown in Fig 2-D, the lowest ΔG corresponded to a kinked binding tunnel. These results contrast with those of the stearoyl-CoA in Substrate-Waterbox model. Over the 100-ns MD, the most frequently observed dihedral was −60° for the stearoyl-CoA in the Substrate model (Table 1-A), whereas it was as opposed to −180° for the stearoyl-CoA in the Substrate-Waterbox model (Table 1-G; Fig 3). In the kinked binding tunnel arrangement, the tunnel was kinked due to the proximity of the residues and the dihedral about C9-C10 was near a reactive position. In this way, SCD1 stabilizes the substrate for desaturation.

The substrate-protein conformation is distinctly different from the product-protein conformation (Scheme 1 step 3 vs. step 4). For example, Substrate and Product models shared similar (within 0.1 Å) paired distances between Gln147-Trp153 and Gln147-Thr261 in only 4 of the 18000 observations. The Substrate model and Apoprotein model were more similar in this respect, including ΔG distribution overlap at lower kT (Fig 2-A,C). In addition to resembling the Substrate model (Table 1-A) in this way, the Apoprotein model (Table 1-C) ΔG distribution overlapped with the Protein model (Fig 2-B,C). These findings indicate that when the protein does not have either a substrate or product present, the two paired distances of residues of interest are free to sample both conformations. Consequently, it is the respective interactions between the saturated stearoyl-CoA substrate or the desaturated oleoyl-CoA with the protein that results in these distinct ΔG distributions.

Consistent with the close proximity of Gln147-Trp153 and Gln147-Thr261 observed in the human and murine crystal structures, the Substrate model (Table 1-A) results indicate that the most energetically favorable conformation was Gln147- Trp153 = 1.95Å, Gln147-Thr261 2.15Å (Fig 2-A). These distances suggest possible hydrogen bonding between the residues and correspond to a kinked tunnel shape (Fig 2-D). On the other hand, the Product model exhibited larger distances between these residues at the ΔG distribution minimum, Gln147- Trp153 = 4.05Å, Gln147-Thr261 = 5.25Å (Fig 2-A). This geometry, inconsistent with hydrogen bonding, corresponds to an open tunnel shape. Taken together, substrate:protein and product:protein interactions are largely different, suggesting a route for substrate-product discrimination related to the positions Trp153-Gln147-Thr261.

While the naturally-occurring substrate had the most frequently observed negative gauche dihedrals about the C9-C10 double bond, there was a substantially higher ratio of negative gauche dihedrals to trans dihedrals than when without the CoA moiety (Table 1-D, E, respectively). While in this way there is a penalty for inclusion of the CoA moiety for the Saturated Lipid (i.e. more unfavorable trans dihedrals), results from the Desaturated Lipid model indicate that substrate-product differentiation is compromised when the CoA moiety is not present (Table 1-D, E respectively). In the naturally occurring Product model (Table 1-B), when there is a double bond between the carbons, these residues, along with the loop at the entrance of the tunnel, move such that the entrance is open, perhaps so the product may exit SCD1. This finding is consistent with the suggestion of Bai *et al*.[19] that in order for the product to leave the tunnel and move into the membrane, the putative hydrogen bond between Gln143-Thr257 in the murine model (equivalent to human SCD1 residues Gln147-Thr261) may be broken, allowing for product egress into the membrane.

## Conclusion

The current results describe the dynamics of SCD1 in the apoprotein state, with interacting CoA headgroup, with bound substrate, and with bound product. This work describes a structure-function relationship characterization of D9D. These findings assist in prioritizing experimental directed mutagenesis work. A sequence of dynamics events drive the protein:substrate recognition and processing and the release of the substrate. As summarized in Scheme 1, results indicate that 1) the Apoprotein has a shallow entrance to the tunnel, 2) the CoA headgroup opens the tunnel entrance, 3) hydrogen bonding between Gln147-Trp153 and Gln147-Thr261 contributes to the tunnel kink typical of the Substrate model, 4) distance between these three residues in the Product model may aid in product release into the membrane. Furthermore, the CoA headgroup may aid in substrate-product discrimination.

## ASSOCIATED CONTENT

### Supporting Information

The SI file includes: SI Figs 1-6, SI Table 1, and link to data repository.

## AUTHOR INFORMATION

### Corresponding Author

*Dr. Jerome Baudry.

### Present Addresses

Same as affiliation.

### Author Contributions

The manuscript was written through contributions of all authors. All authors have given approval to the final version of the manuscript. A.B.P. contributed to study and project design, performed the calculations, and prepared the manuscript. R.L.W. performed initial project calculations and contributed to study design. C.R.Y. and J.D.N. contributed to project design and manuscript preparation. J.B. led study and project design and manuscript preparation.

## ACKNOWLEDGMENT

We thank Edward Chaum and Noriko Inoguchi for helpful discussions.

## ABBREVIATIONS

SCD1: Stearoyl-CoA Desaturase-1
D9D: Delta-9 Desaturase
MUFA: monounsaturated fatty acids
CGenFF: CHARMM General Force Field
POPC: 1-palmitoyl-2-oleoyl-*sn*-glycero-3-phosphocholine
MD: molecular dynamics
MOE: Molecular Operating Environment.

